# Positive genetic associations among fitness traits support evolvability of a reef-building coral under multiple stressors

**DOI:** 10.1101/572321

**Authors:** Rachel M. Wright, Hanaka Mera, Carly D. Kenkel, Maria Nayfa, Line K. Bay, Mikhail V. Matz

## Abstract

Climate change threatens organisms in a variety of interactive ways that requires simultaneous adaptation of multiple traits. Predicting evolutionary responses requires an understanding of the potential for synergistic interactions among stressors and the genetic variance and covariance among fitness-related traits that may reinforce or constrain an adaptive response. Here we investigate the capacity of *Acropora millepora*, a reef-building coral, to adapt to multiple environmental stressors: rising sea surface temperature, ocean acidification, and increased prevalence of infectious diseases. We measured growth rates (weight gain), coral color (a proxy for Symbiodiniaceae density), and survival, in addition to nine physiological indicators of coral and algal health in 40 coral genets exposed to each of these three stressors singly and combined. Individual stressors resulted in predicted responses (*e.g*., corals developed lesions after bacterial challenge and bleached under thermal stress). However, corals did not suffer substantially more when all three stressors were combined. Nor were tradeoffs observed between tolerances to different stressors; instead, individuals performing well under one stressor also tended to perform well under every other stressor. An analysis of genetic correlations between traits revealed positive co-variances, suggesting that selection to multiple stressors will reinforce rather than constrain the simultaneous evolution of traits related to holobiont health (*e.g.,* weight gain and algal density). These findings support the potential for rapid coral adaptation under climate change and emphasize the importance of accounting for corals’ adaptive capacity when predicting the future of coral reefs.

## Introduction

Reef-building corals are experiencing unprecedented declines due to changing environmental conditions, such as rising sea surface temperatures that lead to coral bleaching, and ocean acidification that impairs calcification (Andersson & Gledhill, 2012; Hoegh-Guldberg et al., 2007). Climate change has also indirectly led to increasingly prevalent coral diseases, which are often attributed to the increased abundance and virulence of bacterial pathogens (Ben-Haim, 2003; Maynard et al., 2015; Pruzzo et al., 2010). In the face of these stressors, corals are left with few options but to move, adapt, or die. A number of studies have documented corals’ capacities to expand their ranges to more suitable habitats (Makino et al., 2014; Yamano, Sugihara, & Nomura, 2011). Models that simulate future coral cover under different climactic scenarios increasingly include estimates of adaptive capacity, such as simulated directional genetic selection (Logan, Dunne, Eakin, & Donner, 2014) or predictions of the spread and persistence of alleles conferring thermal-tolerance (Bay, Rose, Logan, & Palumbi, 2017; Matz, Treml, Aglyamova, & Bay, 2018). These studies primarily focus on a single environmental challenge (thermal stress) and do not predict interactive effects among simultaneous stressors or account for genetic associations between multiple tolerance traits.

Adaptation to rapidly changing conditions requires standing phenotypic variation upon which selection can act, provided that this heterogeneity has a genetic basis (*i.e*., that it is heritable). Although a large and ever-growing number of studies examine mean responses of coral species to individual effects of climate change (Marubini, Ferrier-Pages, & Cuif, 2003; Okazaki et al., 2017), few have measured standing genetic variation and heritability of these responses (Dixon et al., 2015; Kenkel, Setta, & Matz, 2015; van Oppen, Császár, Berkelmans, Ralph, & Frankham, 2010; Vollmer & Kline, 2008; S Wang et al., 2009) and even fewer have assessed variation in multiple stress tolerance phenotypes (Shaw, Carpenter, Lantz, & Edmunds, 2016). Corals exhibit remarkable variation in stress tolerance traits upon which selection could theoretically act (Baums et al., 2013; Dixon et al., 2015; Wright et al., 2017). However, univariate analyses assessing a single stress-response phenotype, such as mortality under bacterial challenge or bleaching under thermal stress, fail to fully describe the genetic basis of the phenotypes under selection. Selection is an inherently multivariate process that acts simultaneously on sets of functionally related traits. Indeed, centuries of animal and plant breeding have demonstrated that selection on one trait will often result in changes in another correlated trait (Rauw, Kanis, Noordhuizen-Stassen, & Grommers, 1998; Zhao, Atlin, Bastiaans, & Spiertz, 2006). Commercial demand for multiple-stress-tolerant crops has driven extensive research on stressor combinations in plants (Pandey, Irulappan, Bagavathiannan, & Senthil-Kumar, 2017). In these plants, combinations of abiotic stressors and pathogens result in either resistance or susceptibility to disease, depending on the intensity or duration of stress (Pandey, Ramegowda, & Senthil-Kumar, 2015). Positive associations between different stressors can be attributed to shared pathways. For example, some biotic and abiotic stressors stimulate the same defense-related endogenous signals (Mithöfer, Schulze, & Boland, 2004). Alternatively, stress responses may compete for demands on energetic reserves, resulting in a negative association in tolerances, or a trade-off (Sokolova, 2013).

The prospects for future reef-building corals are exceedingly pessimistic without rapid adaptation to a number of simultaneous stressors. This capacity for adaptation is determined by the answer to an outstanding question: does success under one type of stress come at a cost of susceptibility to a co-occurring environmental challenge? To address this critical knowledge gap, we quantified the capacity for *Acropora millepora*, a model representative of a keystone group of marine organisms that are among the most vulnerable to climate change (Reusch, 2014), to adapt to simultaneous stressors. Multiple coral colonies (n = 40, hereafter referred to as “genets”) were split into replicate clonal fragments (n = 5 per treatment) that were exposed to elevated temperature (30°C), increased *p*CO_2_ (*p*H = 7.8, 700 ppm CO_2_), bacterial challenge (10^6^ CFU mL^-1^ *Vibrio owensii*), a combination of these three stressors at the same levels, or a control condition (27°C, *p*H = 8.0, 400 ppm CO_2_, no added *V. owensii*). We measured a comprehensive suite of coral host and algal traits to assess each genet’s performance in each condition and constructed a genetic variance–covariance matrix to identify potential genetic trade-offs or reinforcements between phenotypes.

## Methods and Materials

### Study organism and aquarium conditions

Forty-one colonies of *Acropora millepora* were sampled between October – December 2014 from Davies Reef lagoon (78 km offshore; 18°50’11’’S, 147°38’41’’E), Rib Reef (56 km offshore; 18°28’55’’S, 146°52’15’’E), Pandora Reef (16 km offshore; 18°48’44’’S, 146°25’59’’E), and Esk Island (24 km offshore; 18°46’04’’S, 146°30’57’’E). These colonies were transferred to holding tanks at the National Sea Simulator system at the Australian Institute of Marine Science (Townsville, Queensland, Australia). After approximately two weeks acclimatization, each colony was fragmented into 25 replicate fragments (“nubbins”), which were mounted on aragonite plugs and placed on replicate trays. Trays were maintained in six indoor holding tanks which were supplied with 0.2 µM filtered seawater (FSW) at 27°C. Three lights (AI Aqua Illumination, USA) were suspended above each tank providing an average underwater light intensity of 180 µmol photons m^-2^ s^-1^ on a 10-/14-hour light–dark cycle. Corals were fed freshly hatched *Artemia nauplii* twice daily and cleaned three times a week to prevent algal growth. Coral nubbins were acclimated to these conditions for 3–5 months, depending on the date of collection. Unique genets were later confirmed via 2b-RAD genotyping (Shi Wang, Meyer, Mckay, & Matz, 2012). The final total genets for each sampling location are as follows: Davies (n = 10), Rib (n = 10), Pandora (n = 14), and Esk (n = 6). Algal symbiont types were investigated by ITS-2 sequencing for eight of the coral colonies (Howe et al., submitted).

### Experimental treatments and sample preparation

On 2 March 2015, coral nubbins (25 per genet) were placed into 25 50 L treatment tanks fitted with 3.5-W Turbelle nanostream 6015 pumps (Tunze, Germany) with flow through seawater at ∼25 L hour^-1^ at the same temperature and light conditions as in the previous holding tanks. Initial weights for each nubbin were obtained following the method described by (Spencer Davies, 1989). Tanks (n = 5 per treatment) were allocated to the following treatments: elevated temperature (30°C), increased *p*CO_2_ (700 ppm, pH = 7.8), bacterial challenge (10^6^ CFU mL^-1^ *Vibrio owensii* DY05), a combined treatment (30°C, 700 ppm, 10^6^ CFU mL^-1^ *V. owensii*), and control (27°C, 400 ppm, pH = 8.0, no bacteria). This isolate of non-quarantined *V. owensii* had been recently sampled during an infectious disease outbreak in cultured lobsters at the research facility. Temperature and *p*CO_2_ were gradually increased in their respective tanks over the course of a week to 30°C and 400 ppm (pH = 8.0). The bacterial challenge was conducted in separate isolated tanks (no flow through) and consisted of a daily six-hour incubation. *Vibrio owensii* was added at a final concentration of 10^6^ CFU mL^-1^ to every bacterial challenge tank, including the combined treatment (which was also maintained at 30°C and 700 ppm *p*CO_2_ for these six hours). Corals were then returned to their respective treatment tanks until the next day’s bacterial challenge and the bacterial challenge tanks were treated with 20% bleach for at least 30 minutes. Coral fragments were photographed daily to quantify bleaching via the Coral Health Chart (Siebeck, Marshall, Klüter, & Hoegh-Guldberg, 2006) and lesion progression. Net oxygen production and the change total alkalinity under light were measured for randomly selected genets following methods described in (Strahl et al., 2015). Fragments exhibiting any tissue loss or that were fully bleached and exhibiting algal growth were removed from treatment tanks, buoyant weighed, and preserved in liquid nitrogen. The time of death (day post initial exposure) was recorded at each instance. The experiment continued for 10 days, until approximately 21% mortality was recorded over all treatments. Eleven days after the initial challenge, all surviving corals were photographed, buoyant weighed, preserved in liquid nitrogen, and stored at −80°C until sample processing.

Tissue was removed from coral skeletons using an air gun and 0.2 µM filtered seawater and homogenized for 60 seconds using a Pro250 homogenizer (Perth Scientific Equipment, Australia). A 1 mL aliquot of the tissue homogenate was centrifuged for 3 minutes at 1500 ×*g* at 4°C and the pellet was stored at −80°C for chlorophyll analyses. The remaining homogenate was centrifuged for 3 minutes at 1500 ×*g* at 4°C to separate host and symbiont fractions. The fractions were frozen in 96-well tissue culture plates and stored at −80°C. Coral skeletons were rinsed with 10% bleach then dried at room temperature (∼24°C). Skeletal surface area was quantified using the single paraffin wax dipping method (Stimson & Kinzie, 1991) and skeletal volume was determined by calculating water displacement in a graduated cylinder.

### Physiological trait assays

Assays were conducted to detect cellular and metabolic activity changes within the Symbiodiniaceae or coral host tissue in response to the treatment. All standards and samples were loaded as duplicates, and absorbance was recorded with a Cytation 3 multi-mode microplate reader (BioTek, Winooski, USA) and analyzed using Gen5 software (BioTek, Winooski, USA).

To quantify chlorophyll concentrations, tissue homogenate algal pellets were resuspended in 1 mL chilled 90% acetone. The homogenate was sonicated on ice for 10 seconds at 40% amplitude, left in the dark for 20 minutes, and centrifuged for 5 minutes at 10,000 ×*g* at 4°C. A 200 µL aliquot of sample extract was loaded to a 96-well plate, and absorbance was recorded at 630 and 663 nm. Chlorophyll a and c2 concentrations were calculated with the equations in Jeffrey & Haxo (1968) and were normalized to surface area:

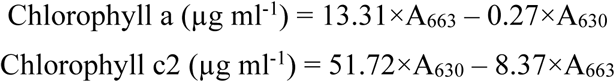

A commercial colorimetric protein assay kit (DCTM Protein Assay Kit, Bio-Rad, Hercules, USA) was used to quantify total protein content of the coral host tissue. A 100 µL aliquot of Symbiodiniaceae-free coral tissue sample was digested using 100 µL sodium hydroxide in a 96-well plate for 1 hour at 90°C. The plate was centrifuged for 3 minutes at 1500 ×*g*. Following manufacturer’s instructions, 5 µL digested tissue was mixed with 25 µL alkaline copper tartrate solution and 200 µL dilute Folin reagent in a fresh 96-well plate. Absorbance at 750 nm was recorded after a 15-minute incubation. Sample protein concentrations were calculated using a standard curve of bovine serum albumin ranging from 0 and 1000 µg mL^-1^.

Carbohydrate content of the Symbiodiniaceae-free coral tissue was estimated following the method by Masuko et al. (2005) that measures monosaccharides, including glucose, which is the major photosynthate translocated between symbionts and host corals (Burriesci, Raab, & Pringle, 2012). A 50 µL aliquot of coral tissue was mixed with 150 µL concentrated sulfuric acid and 30 µL 5% phenol in a 96-well plate for 5 minutes at 90°C. After another 5-minute incubation at room temperature, absorbance at 485 nm was recorded. The total carbohydrate concentrations of samples were calculated using a standard curve of D-glucose solutions ranging from 0 to 2000 µg ml^-1^.

To analyze non-fluorescent chromoprotein content, a 30 µL aliquot of coral tissue was loaded to a black/clear 384-well plate and absorbance was recorded at 588 nm. Mean absorbance was standardized to sample protein content.

The activity of catalase (CA), a reactive oxygen species scavenging enzyme (Lesser, 2006), was measured by estimating the change in hydrogen peroxide (H_2_O_2_) substrate concentration. A 20 µL aliquot of coral tissue was mixed with 30 µL 50 mM phosphate-buffered saline solution (PBS; pH 7.0) and 50 µL 50 mM H_2_O_2_ in a 96-well plate. CA was calculated as the change in absorbance at 240 nm every 30 seconds over the linear portion of the reaction curve for 15 minutes and was standardized to sample protein content:

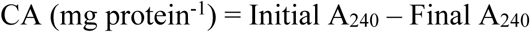

The change in coral color was estimated using photographs taken during the experiment with a Nikon D300 digital camera. Brightness values were measured over the entire front and back sections of each nubbin using image analysis software (ImageJ, NIH). Corals become brighter (whiter) when they lose darkly colored algal symbionts, so changes in brightness reflect changes in Symbiodiniaceae densities (Beer, Loya, Winters, Holzman, & Blekhman, 2009). A standard curve of brightness values was constructed using standard coral color cards that were present in each image. Brightness values were standardized to color cards to normalize for differences across photo sessions.

### Statistical analyses

The time of death was noted for each individual nubbin. Survival was modeled as time of death ∼ treat + reef + (1|tank) using *coxme* package in R 3.3.1 (R Core Team, 2016; Therneau, 2012), where treatment was specified as the presence or absence of elevated heat, bacteria, or increased *p*CO_2_ (*i.e*., bac = 1 for “bacteria” and “all” treatments; heat = 1 for “heat” and “all” treatments). All subsequent analyses were performed on lesion-free (alive) corals. Data were log-transformed using powers determined by Box-Cox transformations performed using the *powerTransform* function in the R package *car* (Fox & Weisberg, 2001). Linear mixed-effects models implemented using the R package *nlme* (Pinheiro, Bates, DebRoy, & Sarkar, 2017) tested the effects of treatments, treatment interactions, and reef-of-origin on trait values. The *stepAIC* function in the R package *MASS* (Venables & Ripley, 2002) determined which terms to include in the best-fit model. Principal components analysis on mean-centered and variance-scaled values was performed using the *prcomp* function in base R. Pearson correlations between trait values were calculated using the cor function in base R, and correlation heatmaps were constructed using the *corrplot* function. The genetic variance–covariance matrix was constructed using the R package MCMCglmm (Hadfield, 2010). Trait data were mean-centered and variance scaled. The multivariate model was fit for four traits (growth, color, chlorophyll c2, and carbohydrate) with treatment as a fixed effect and genet as a random effect, using the idh variance structure. The model was run for 20000 iterations after a 5000 iteration burn-in, storing the Markov chain after 20 iteration intervals. Partial regression coefficients for each trait on binomial survival outcome were modeled using a categorical MCMCglmm model with genet as a random effect. The selection gradient was composed of these partial regression coefficients, scaled to unit variance. Predicted changes in trait values 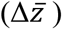 were calculated using the multivariate breeder’s equation (Lande & Arnold, 1983):

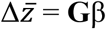

**G** is the genetic variance–covariance matrix and β is the selection gradient.

## Results

### Mean responses to treatments

#### Survival

Corals in the control condition experienced the lowest mortality at 13.5%, followed by increased temperature (21.5%), combined treatment (23.7%), bacteria challenge (26.4%), and elevated *p*CO_2_ (27.4%). Mortality rates were compared using the *coxme* package in R (R Core Team, 2016; Therneau, 2012). Treatment was specified as the presence or absence of each stressor. The Cox proportional hazards model included each treatment, interactions among treatments, and reef as fixed effects, with tank as a random effect. Bacteria challenge significantly increased mortality rates (hazard ratio [HR] = 3.32, p = 0.018), while a weak interaction between bacteria challenge and increased temperature slightly improved survival odds (HR = 0.17; p = 0.09). Elevated *p*CO_2_ ultimately caused the most mortality, but mortality rates in corals under this condition were indistinguishable from control corals throughout the first week of the experiment (Figure 1A). Colonies from Rib reef had the lowest mortality (16.5%). Colonies from Davies and Esk reefs had higher mortality rates (HR = 0.54, p = 0.007 and HR = 0.58, p = 0.01, respectively) than those from Rib (Supplementary Figure 1).

**Figure 1:**
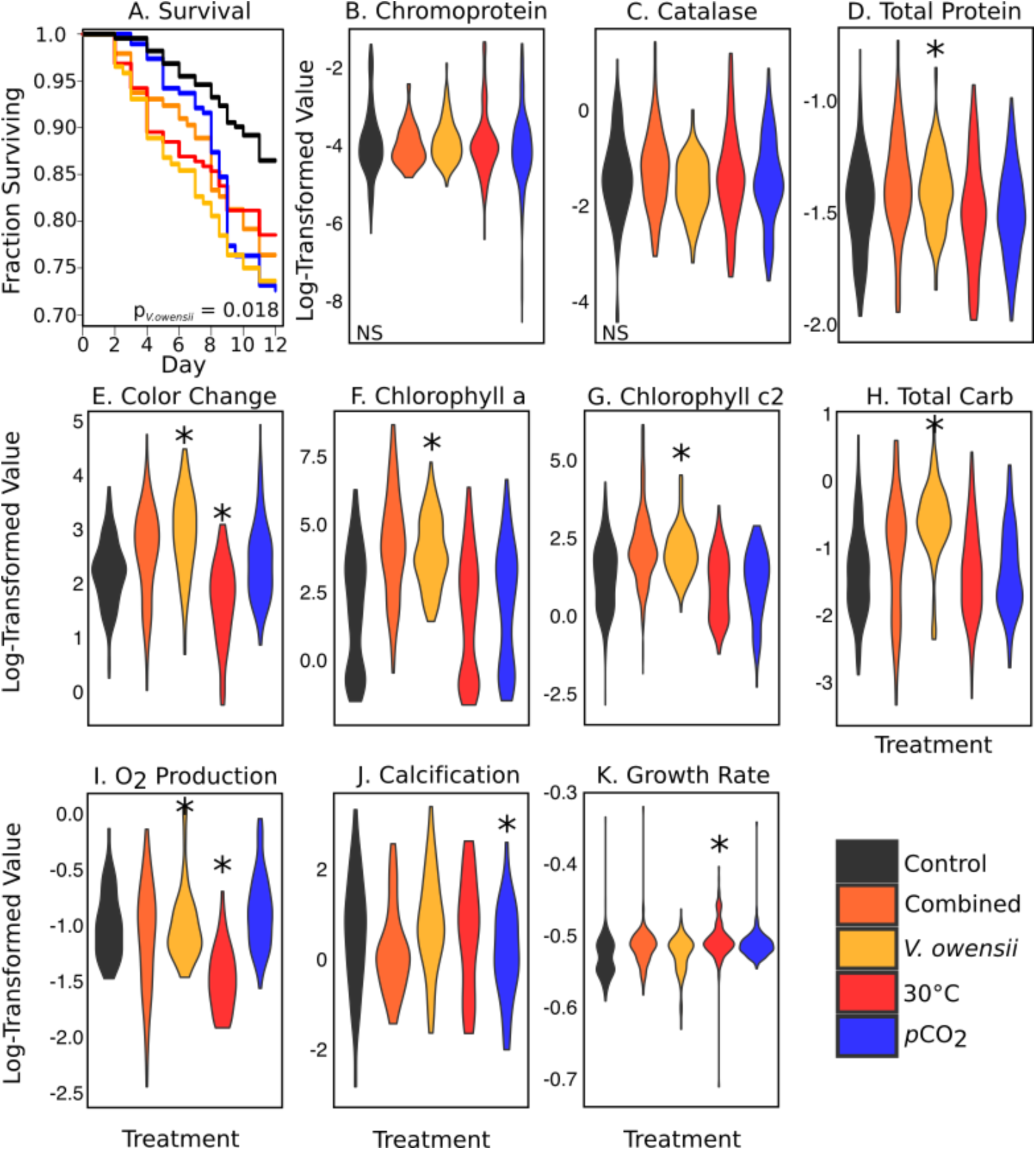
Mean responses to treatment. (A) Survival fraction over the duration of the experiment. Colors correspond to treatment (see inset key). (B–K) Box-Cox power transformed trait values separated by treatment (indicated by color in legend). NS = no significant effect of treatment. Asterisks indicate a significant (p < 0.05) effect of the indicated treatment. (B) Chromoprotein content (A_588_ · µg protein^-1^). (C) Catalase activity (?H_2_O_2_ · mg protein^-1^ · min^-1^). (D) Total protein content (mg · cm^-2^). (E) Coral fragment color change (final–initial score). Chlorophyll a (F) and c2 (G) content (µg · cm^-2^). (H) Total carbohydrate (mg · cm^-2^). (I) Oxygen production (µg O_2_ · cm^-2^ · min^-1^). (J) Instant calcification rate (µmol CaCO_3_ · cm^-2^ · min^-1^). (K) Buoyant weight growth rate (% Δ weight g · day^-1^).

#### Physiological Responses

We measured the following algal and host-associated traits from surviving coral fragments: coral color (an indicator of Symbiodiniaceae densities), algal chlorophyll a and chlorophyll c2 content, total host carbohydrate, total host protein, host catalase activity, host chromoprotein content, oxygen production (indicator of photosynthetic rate), instant calcification rate, and change in buoyant weight (skeletal growth).

Chromoprotein content and catalase (Figure 1B–C) serve as proxies for coral innate immune response. We found no significant effect of treatment on either of these measurements. However, there was a weak interaction between the effects of elevated temperature and bacterial treatment, which tended to increase catalase activity (β = 0.33, p = 0.06).

Bleaching was calculated as a log-transformed change in color in photographs standardized to the Coral Health Chart (Siebeck et al., 2006). Bacteria treatment made corals significantly darker (β = 0.70, p < 0.001), whereas corals became lighter (*i.e.,* bleached) in the elevated temperature treatment (β = −0.54, p < 0.001; Figure 1E). These findings are corroborated by chlorophyll measurements: bacterial treatment increased chlorophyll a (β = 2.1; p < 0.001; Figure 1F) and c2 (β = 1.0, p = 0.001; Figure 1G) content in the algal fraction of the coral tissue. Bacteria treatment increased the carbohydrate content in the coral host tissues (β = 0.73, p < 0.001; Figure 1H). We also observed a weak negative interaction between elevated temperature and bacterial challenge on carbohydrate content (β = −0.54, p = 0.059).

Photosynthetic and instant calcification rates were measured for a smaller subset of the coral genets. As expected, the elevated temperatures reduced photosynthetic rates (β = −0.51, p < 0.001, Figure 1I). Coincident with improved algal traits under bacterial challenge (Figure 1E–G), the bacterial treatment rescued photosynthetic rates under elevated temperatures and there was a positive interaction between these two stressors (β = 0.40, p = 0.04; Figure 1I). Only the elevated *p*CO_2_ treatment affected instant calcification rates, decreasing them on average (β = −0.48, p = 0.01; Figure 1J). Buoyant weights of each fragment were measured at the beginning of the experiment and when each fragment was removed from the experiment. Corals in the elevated temperature treatment experienced moderately increased growth rates (β = 0.017, p = 0.04, Figure 1K). Buoyant weights were not significantly affected by any other treatment.

### Phenotypic space, correlations, and evolvability calculations

The lack of synergistic treatment effects on coral fitness proxies provides encouraging evidence for an individual coral’s capacity to resist multiple stressors. To investigate whether a population of corals can adapt to multiple threats, we looked for potential tradeoffs by measuring correlations between stressor effects across genets.

Principal components analysis explored patterns in phenotypic space of 429 individual fragments (Figure 2A) with complete datasets for growth, color change, chlorophyll a, chlorophyll c2, chromoprotein, catalase, and survival fraction (the proportion of fragments surviving for the genet in each respective treatment). The first principal component explained 30.2% of the variation and separates samples by differential algal responses to treatment (Figure 2A–B). The second PC explained 16.2 % of the variation and separates samples by host immune enzyme responses (chromoprotein and catalase activity). Holobiont fitness metrics (growth and survival fraction) are projected in the third PC, which represents about 13% of the total variance (Figure 2C).

**Figure 2:**
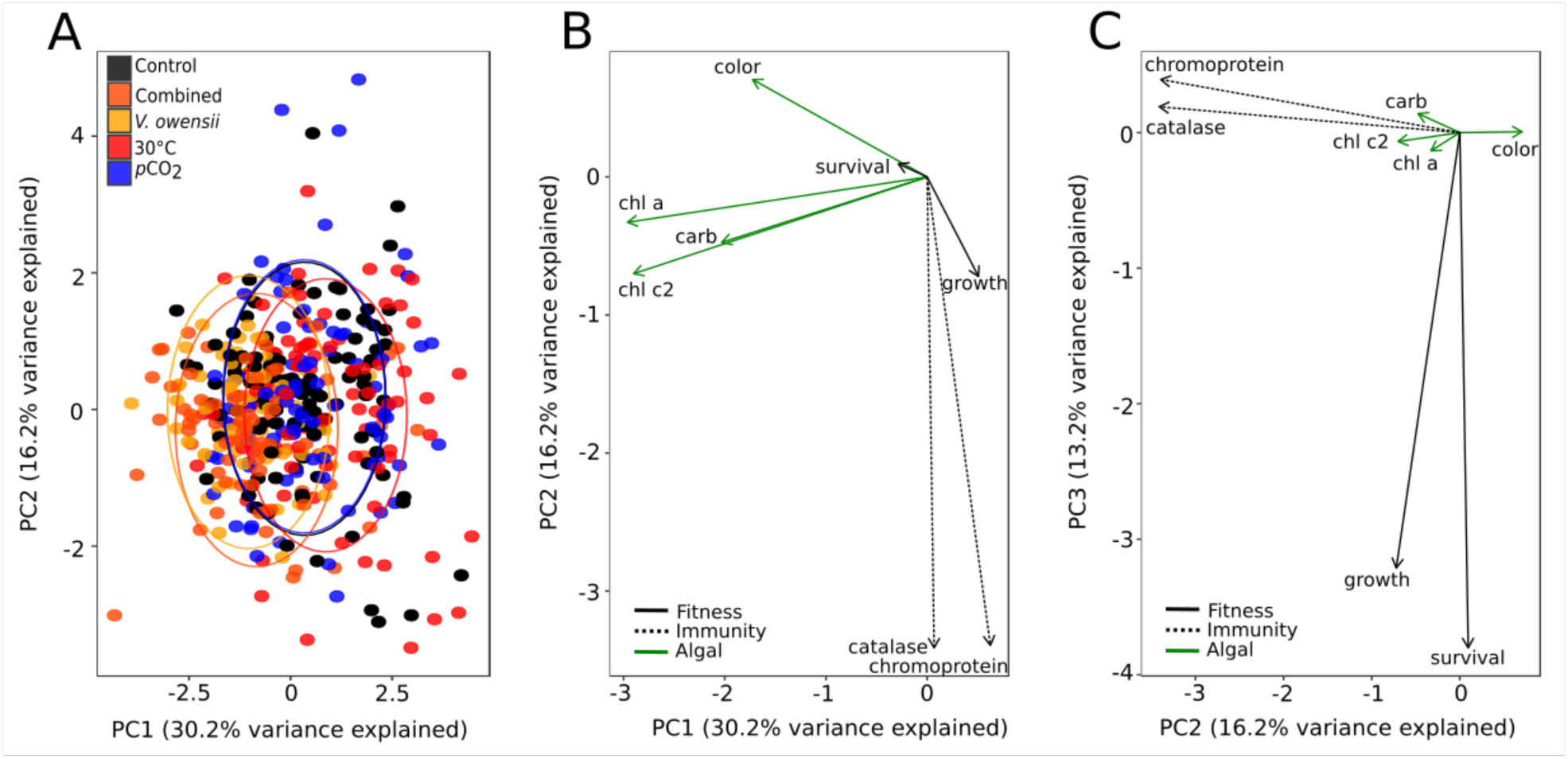
Principal components analysis. (A) Principal components analysis based on physiological trait data for 429 fragments. (B) Loadings for traits along the first two (B) and second and third (C) principal components axes. Data point colors represent the treatment in which the coral fragment was placed. Colors and line shapes of loadings identify traits related to algal parameters (green), immune enzyme activity (dashed), or general coral fitness (black).

We calculated Pearson correlations by genet across treatments for five traits with the most complete data and clearest link to fitness: survival fraction, growth rate, color change, chlorophyll c2 content, and total host carbohydrate (Figure 3). The correlation heatmaps show many positive and statistically significant correlations between trait pairs and very few negative correlations, none of which were statistically significant (Figure 3). Survival fractions for each genet are significantly positively correlated among treatments (Figure 3A). In other words, individuals that show high fitness characteristics under one stressor also perform well under other stressors.

**Figure 3:**
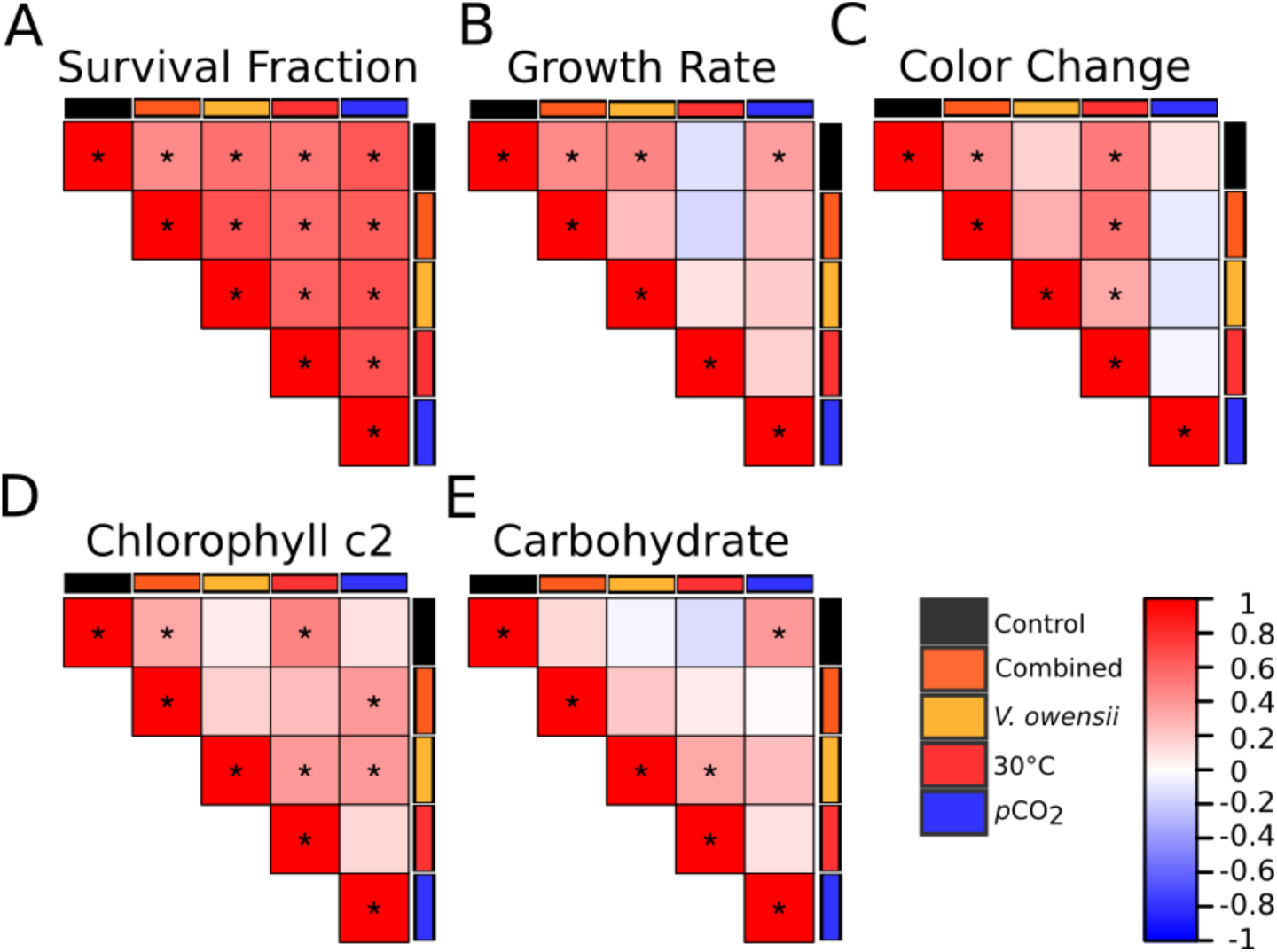
Pearson correlation heatmaps based on scaled average (A) survival fraction, (B) growth rate, (C) color change, (D) chlorophyll c2 content, and (E) carbohydrate content for 39 genets. Colored bars indicate the treatment measurement. Colors within the heatmap squares represent the magnitude and direction of the Pearson correlation according to the key. Significant (p < 0.05) correlations are indicated with asterisks.

Pairwise comparisons do not adequately capture the overall genetic covariance of traits in the population. To further explore the evolvability of these traits, we constructed a genetic variance–covariance matrix (**G** matrix) by selecting four traits that describe various aspects of host and symbiont fitness: color change, weight gain, chlorophyll c2 content, and carbohydrate content. All genetic correlations between these traits were positive (Figure 4A).

**Figure 4:**
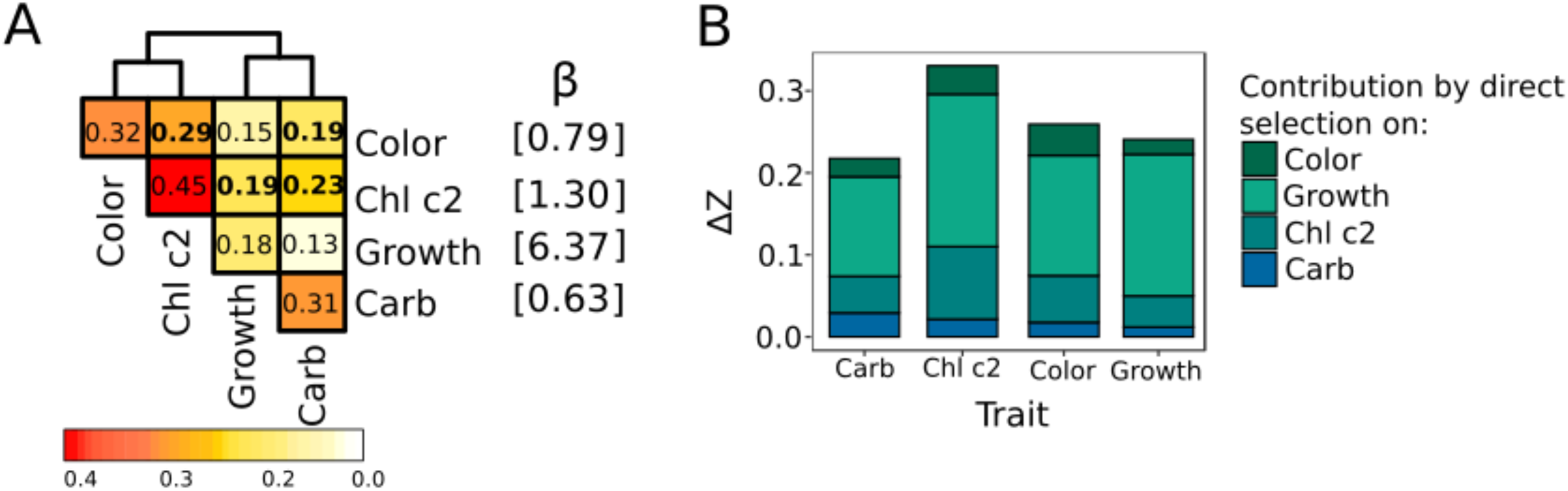
(A) Left: Genetic variance–covariance matrix for four fitness traits across five treatments in 39 genets. The color in the inset key and number within each box quantifies trait variances (diagonal elements) and covariances between paired traits (off-diagonal elements). Bold font denotes significant associations. Right: Selection gradient (β), a vector of partial regression coefficients for each trait on survival. (B) Changes in trait means (ΔZ) per unit of selection for higher survival under multiple stressors. Each stacked bar is composed of the direct effect of selection on the trait and effects of selection on each of the genetically correlated traits.

In multivariate trait space, the response of a trait to selection may deviate from the direction of selection due to influences of genetically associated traits. Depending on the shape of genetic variance–covariance matrix (**G** matrix) and selection strength on individual traits, fitness traits may or may not be able to evolve in concert. To investigate this issue in our coral, we calculated a selection gradient (a vector of partial regression coefficients standardized to unit length) by regressing the four traits against binomial survival (1 = fragment survived; 0 = fragment died). All selection coefficients were positive (Figure 4A), although only growth was significantly associated with survival (p < 0.001). We then applied the multivariate breeder’s equation to estimate how trait values would change given our **G** matrix and selection for higher survival under moderate stressors (Figure 4B). Since all selection coefficients and covariances were positive, the change in every trait over one generation was also positive (Figure 4B). This result implies that fitness traits should be able to co-evolve together and moreover, reinforce each other’s evolution, *i.e*., corals with high survival rates would also tend to produce offspring with increased growth rates, carbohydrate content, Symbiodiniaceae densities, and chlorophyll content under different stressors.

To demonstrate this reinforcement effect, we decomposed the total predicted change in each trait (the total height of each bar in Figure 4B) into the contribution of selection directly on that trait or on the three covarying traits. For example, a predicted increase in chlorophyll c2 content is mostly driven by selection on growth rate, which is more strongly correlated with survival and is genetically correlated with chlorophyll c2 content (Figure 4A). The model predicts less improvement in color (a proxy of algal symbiont density) and host carbohydrate content because these traits covary less with growth rate and are not themselves strongly associated with survival (Figure 4A).

## Discussion

### Mean effects of single and combined stressors

Each experimental treatment largely resulted in the expected mean response. Elevated temperatures reduced algal parameters (Symbiodiniaceae densities, chlorophyll a and c2 content), analogous to a natural coral bleaching event. Bacterial challenge resulted in the development of lesions, but also caused an unexpected increase in algal traits. Although a meta-analysis of coral stress responses predicted that most stressors act synergistically (Ban, Graham, & Connolly, 2014) and other empirical studies have documented compromised coral immunity and increased prevalence of coral disease concurrent with thermal stress (Palmer, 2018; Selig et al., 2007), we did not observe detrimental interactive effects of the combined challenge. In fact, the combination of treatments tended to improve holobiont health relative to a single stress. For example, bacterial challenge alone caused more mortality than the combined treatment (Figure 1A). A tempting hypothesis to explain this phenomenon is that elevated temperature exposure prior to pathogen exposure primed the coral’s stress response system to mitigate oxidative stress associated with launching an innate immune response (Lu, Wang, & Liu, 2015).

We also saw that coral bleaching was minimized in the combined treatment relative to thermal stress alone (Figure 2A), possibly as a result of heterotrophic feeding (Bourne, Morrow, & Webster, 2016): both coral host and algal symbiont had access to extra nutrients (bacterial inoculations triply washed in sterile seawater) in the combined treatment. Further supporting this hypothesis is the unexpected observation of increased algal traits in the bacteria-only treatment. Reef-building corals feed on bacteria (Houlbrèque & Ferrier-Pagès, 2009) which could act as a beneficial nutrient source to encourage algal productivity (Rädecker, Pogoreutz, Voolstra, Wiedenmann, & Wild, 2015; Sawall, Al-Sofyani, Banguera-Hinestroza, & Voolstra, 2014). Heterotrophic compensation has been investigated as a method by which corals withstand extended bleaching events (Baumann, Grottoli, Hughes, & Matsui, 2014; Grottoli, Rodrigues, & Palardy, 2006; Hughes & Grottoli, 2013), but the capacity for heterotrophic feeding to prevent bleaching deserves more attention.

### Best performing genets tolerate multiple stressors

Pairwise correlations of trait values in each treatment revealed mostly positive associations across all genets. Importantly, survival rates were significantly positively correlated under most treatments (Figure 2 and Figure S2), indicating the absence of tradeoffs with respect to an individual’s ability to withstand different disturbances. Individuals that could survive one stressful condition tended to be able to manage other stressors as well. There were a few exceptions involving Symbiodiniaceae: color under bacterial challenge was negatively correlated with survival under bacterial challenge or under thermal stress, and also negatively correlated with growth under control conditions and under bacterial challenge. Again, Symbiodiniaceae density measurements in corals exposed to *Vibrio* in this experiment may reflect the ability of a coral to improve algal traits due to bacteria presence as a result of increased heterotrophic feeding.

Given the drastically different phenotypic responses to each stressor applied in this study (*e.g*., symbiont expulsion *vs*. tissue loss), it is reasonable to expect to observe tradeoffs in a coral’s ability to manage each challenge. Presumably these unique stressors require the coral to employ distinct molecular response mechanisms, which come at a metabolic cost. This study cannot offer a mechanism to explain the positive associations in stress tolerances observed in these corals, but the literature offers promising areas for further investigation. In corals, the oxidative theory for bleaching under thermal stress posits that ROS accumulation damages cells and triggers symbiont expulsion (Lesser, 1997). ROS are also produced in response to immune challenges to exert antimicrobial activity and stimulate immune signaling pathways (Bogdan, Röllinghoff, & Diefenbach, 2000). Innate immune activation limits pathogen growth but also poses immunopathological risk to the host; thus, a maximal immune response is not always optimal (Viney, Riley, & Buchanan, 2005) and antioxidant mitigation of self-harm is necessary for survival (Knight, 2000). Elevated *p*CO_2_ has also been shown to trigger oxidative stress in corals (Davies, Marchetti, Ries, & Castillo, 2016) and oysters (Tomanek, Zuzow, Ivanina, Beniash, & Sokolova, 2011). Given the critical role of managing oxidative stress in responses to thermal stress, elevated *p*CO_2_, and bacterial challenge, the robustness of a coral’s antioxidant defense system to prevent self-harm may underlie tolerance to all three of these stressors. A recent study in rice identified an allele of the transcription factor Ideal Plant Architecture 1 (IPA1) that simultaneously confers improved growth and immune function by toggling between phosphorylation states that drive expression in distinct subsets of gene targets (Wang et al., 2018). Future studies should critically evaluate the nature of shared pathways in coral stress responses.

Another consideration is the contribution of the algal symbiont towards holobiont health. In this study, colonies primarily hosted *Cladocopium* symbionts almost exclusively), but also contained lower abundances of *Brevolium, Durusdinium*, and/or *Gerakladinium* symbionts (<0.1% of the total community) (Howe et al., submitted). Notably, one colony contained a larger proportion of *Durusdinium* and was among the worst performing colonies in this experiment. Though we do not observe a clear contribution of any one symbiont type on overall host health in this study, as other studies have demonstrated (Rouzé, Lecellier, Saulnier, & Berteaux-Lecellier, 2016; Silverstein, Cunning, & Baker, 2017), future studies should continue to investigate how algal symbionts modify coral responses to environmental stressors.

### Positive genetic associations of fitness traits suggest holobiont adaptability

Multivariate analyses describe how fitness traits may change under directional selection. The variance–covariance structure estimated in this study indicates that our focal species possesses the genetic heterogeneity and flexibility to respond to multiple selective pressures. The positive genetic covariances between four traits associated with holobiont fitness (growth, Symbiodiniaceae density, chlorophyll c2 content, and total carbohydrate content) argue for reinforced evolution of all traits (Figure 4). Growth rate is most strongly associated with survival under climate change stressors and therefore would evolve rapidly due to direct selection pressure. Although directional selection is weaker for other traits (algal density, carbohydrate content, and chlorophyll c2 content), genetic correlation with growth rate would result in their increases as well. However, it is important to note that our experiment cannot disentangle genetic associations that are due to host, symbiont, or their specific combination. Our conclusions regarding the effect of selection on fitness traits would hold only if genetic interactions between host and symbiont do not contribute much to fitness trait variation (*i.e*., if most of the trait variance is attributable to a simple sum of variances due to host and symbiont). More research on study systems where holobiont genetic composition can be manipulated (*e.g*., coral recruits) is necessary to resolve this issue.

### Considerations for future reefs

The management implications of these findings are two-fold. Firstly, adaptive processes should not be ignored in ecological climate modeling. Dire estimates for future coral cover are often derived from experiments in which a coral from today is placed under conditions predicted decades into the future (Okazaki et al., 2017). *A. millepora* can reach reproductive maturity as early as three-years after fertilization (Baria, dela Cruz, Villanueva, & Guest, 2012) and thus, modeling strategies based on end-of-the-century climate scenarios ignore dozens of generations of potential adaptive evolution. Our results suggest that this adaptation can proceed because coral fitness traits tend to reinforce, rather than constrain, adaptation toward improved fitness under multiple environmental challenges. Secondly, our results should be taken into account during efforts to spread adaptive genetic variation by propagating, translocating, and breeding genets that have survived a natural stress event, although latent effects of the stress event may impact thermal tolerance into the future. Regardless of the method used to select broodstock for coral reseeding, our results strongly suggest that colonies should be selected for restoration in a manner that does not jeopardize corals’ natural ability to adapt, for example through a narrowing of the gene pool. In the absence of severe bottlenecks in genetic diversity, natural selection will continue selecting for corals that thrive despite multiple harassments brought about by climate change. However, this adaptation will only proceed if reproduction is maintained under increasingly hostile conditions and until adaptive genetic variation starts running out (Matz et al., 2018), and therefore our findings should not undermine the critical urgency to limit anthropogenic climate change.

## Acknowledgements

We would like to thank the staff of AIMS research vessel the RV Cape Ferguson and the National Sea Simulator for assistance with coral colony collection and experiments. This work was supported by funding provided by AIMS to LBK and CDK, NSF DBI-1401165 to CDK. The collections were supported by a permit from the Great Barrier Reef Marine Park Authority permit numbers G11/34671.1 and G14/37318.1.

