## Supplemental Figures for "Positive genetic associations among fitness traits support evolvability of a reef-building coral under multiple stressors"

**Supplementary Figures**

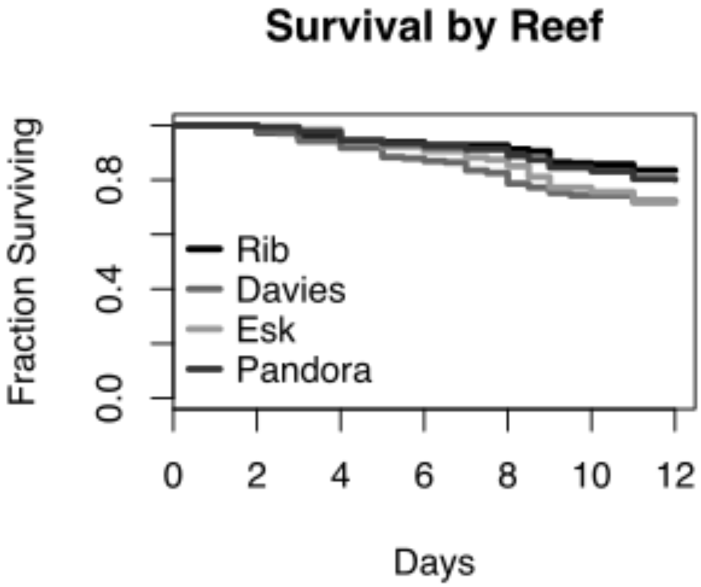


Supplementary Figure 1: Survival rates by reef of origin.

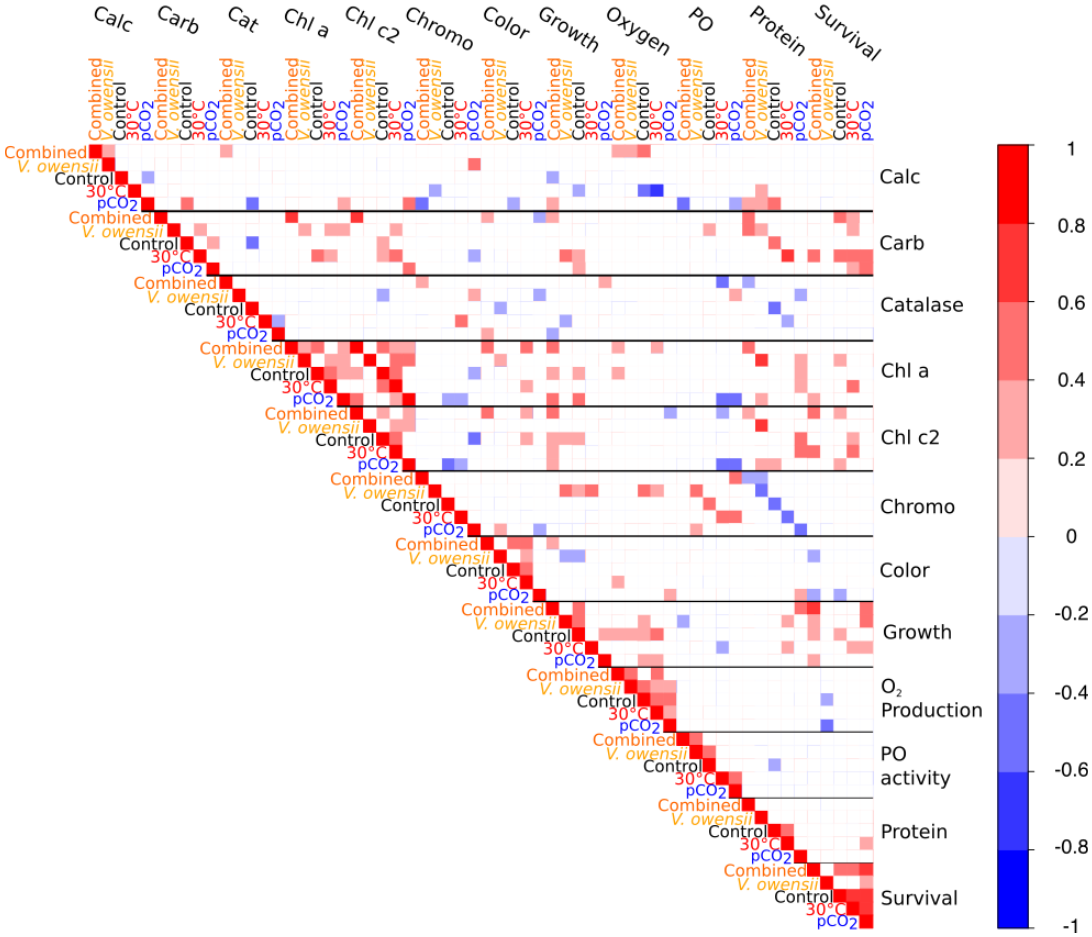


Supplementary Figure 2: Pearson correlation heatmap based on scaled average phenotypic values for 38 genotypes. Labels indicate the treatment and trait measurement. Colors within the heatmap squares represent the magnitude and direction of the Pearson correlation according to the inset key. Only significant ($p < 0.05$) correlations are shown.
